# Prophylactic and therapeutic antiviral effects of the influenza A defective interfering particle OP7 in human lung epithelial cells *in vitro*

**DOI:** 10.64898/2026.03.30.715239

**Authors:** Patricia Opitz, Jan Küchler, Kristina Holdt, Elias Hofmann, Daniel Rüdiger, Sascha Y. Kupke, Udo Reichl

**Affiliations:** Bioprocess Engineering, Otto von Guericke University, Magdeburg, Germany; Bioprocess Engineering, Max Planck Institute for Dynamics of Complex Technical Systems, Magdeburg, Germany

**Keywords:** Influenza A virus, defective interfering particles, antiviral mechanisms, antiviral prophylaxis, therapeutic interference

## Abstract

Defective interfering particles (DIPs) derived from the influenza A virus (IAV) are a promising antiviral agent due to their strong antiviral efficacy demonstrated in various animal models. OP7 is an unconventional IAV DIP with multiple point mutations in the viral RNA (vRNA) of genome segment 7, as opposed to the large internal genomic deletions typically found in conventional IAV DIPs. Further, OP7 showed an even higher interfering efficacy than conventional DIPs. However, the inhibitory effect of OP7 on standard virus (STV) replication has primarily been investigated in Madin-Darby Canine Kidney (MDCK) cells, which lack a functional myxovirus resistance (Mx)-mediated antiviral activity against IAV. In this study, we examined the antiviral activity and mechanism of antiviral action of OP7 in an interferon (IFN)-competent human lung carcinoma cell line (Calu-3) *in vitro*. We performed STV and OP7 co-infection experiments using a variety of infection conditions and measured the time-resolved dynamics in viral titer, vRNA, protein level, and host cell gene expression. We observed that OP7 co-infection results in enhanced type I IFN responses and markedly reduced infectious virus release, even at low doses. Additionally, we found that at a high STV multiplicity of infection (MOI), the replication interference of OP7, suppressing the replication of STV vRNA, appears to be the dominant mechanism of its antiviral action. At a low MOI, however, IFN induction seems to be more important. Furthermore, we examined the efficacious co-infection time window for potential prophylactic and therapeutic antiviral treatment. We observed an antiviral effect exerted by OP7 infection for up to seven days before STV infection and up to 24 hours after STV infection. Together, these findings demonstrate that OP7 is a potent antiviral DIP. Therefore, this work supports the further development of OP7 as a therapeutic and prophylactic antiviral agent.

## Introduction

The influenza A virus (IAV) poses a threat to global health. It infects one billion people every year, placing a significant strain on healthcare systems (Thompson et al., 2009). Annual prophylactic vaccination against the seasonal strains is the most effective way to prevent infection and the spread of the virus (Gebre et al., 2021). However, the production of vaccines takes several months from strain selection to manufacturing and release (Mokalla et al., 2025). Therefore, antivirals are important for complementing vaccination efforts due to their rapid mode of action. In addition, antiviral use is a vital tool for pandemic preparedness, enabling a rapid response to emerging viral threats. Defective interfering particles (DIPs) are considered promising candidates for antiviral applications and have been the subject of research in recent years (Alnaji & Brooke, 2020; Levi et al., 2024; Lohmann et al., 2025; Ranum et al., 2024).

IAV-derived DIPs have been proposed as antivirals against IAV infections (Dimmock et al., 2008, 2012; Dogra et al., 2023; Hein, Arora, et al., 2021) as well as against infections of unrelated viruses like the influenza B virus, respiratory syncytial virus, SARS-CoV-2, yellow fever virus and Zika virus (Pelz et al., 2023; Rand et al., 2021; Scott et al., 2011a). Several animal studies have demonstrated the potent anti IAV activity of IAV DIPs (Dimmock et al., 2012; Dogra et al., 2023; Hein, Arora, et al., 2021; Hein, Kollmus, et al., 2021; Huo, Tian, et al., 2020; Zhao et al., 2018). Intranasal administration of IAV DIPs was well tolerated in mice (Dogra et al., 2023; Hein, Kollmus, et al., 2021). In addition, mice were rescued from lethal IAV infection, when DIPs were administered simultaneously (Huo, Cheng, et al., 2020; Scott et al., 2011b; Wang et al., 2023). In previous animal studies, treatment windows for the use of DIPs for prophylaxis and therapy have been evaluated. For prophylaxis, DIP administration two to seven days prior to infection demonstrated significant antiviral effects depending on the virus DIP species (Zhao et al., 2018, 2022; Dimmock et al., 2008; Y. Xiao et al., 2021). For therapy, a time window of six hours to two days was identified (Chaturvedi et al., 2021; Dimmock et al., 2008; Y. Xiao et al., 2021; Zhao et al., 2018).

IAV DIPs typically contain a large internal deletion in one of their eight viral RNA (vRNA) segments and can only replicate upon co-infection with infectious standard virus (STV). The STV provides the missing gene function that facilitates replication of DIPs when co-infected. In such a scenario, DIPs heavily impair viral replication (Chaturvedi et al., 2021; Levi et al., 2021; Lin Min-Hsuan et al., 2022; Y. Xiao et al., 2021; Yao et al., 2021). DIPs exert their antiviral activity through two principal mechanisms: replication interference (Bdeir et al., 2021; Lin et al., 2025) and stimulation of the interferon (IFN) system (Killip et al., 2015; Manzoni & López, 2018; Tapia et al., 2013). First, DIPs interfere with viral replication because their defective interfering vRNAs (DI-vRNAs) are shorter and replicate faster than their full-length counterparts (Akkina et al., 1984; Mendes & Russell, 2021). Consequently, DI-vRNAs accumulate to higher levels, and therefore deplete essential viral components such as polymerases and nucleoproteins. Thus, they outcompete full-length segments during replication (Laske et al., 2016; Perrault, 1981; Wu et al., 2022). Second, DIPs are potent inducers of the IFN response (Killip et al., 2015; Manzoni & López, 2018; Pavlovic et al., 1992; Penn et al., 2022; Welch et al., 2022; Y. Xiao et al., 2021; Zhou et al., 2023). In the case of an IAV infection, it has been suggested that the cytosolic sensor Retinoic acid-inducible gene I (RIG-I) recognizes DI-vRNAs produced by DIPs, leading to the robust activation of the IFN signaling cascade and the induction of interferon-stimulated genes (ISGs) (Baum et al., 2010; Sengupta & Chattopadhyay, 2024). This results in the establishment of a strong antiviral state in DIP-infected and STV/DIP co-infected cells. However, the relative contribution of replication interference and IFN induction to the antiviral activity of IAV DIPs remains unclear.

Type I IFNs play a central role in the cellular antiviral defense. They bind to their cognate Interferon-α/β receptors and activate Janus kinase 1/2 (JAK1/2), which activates the JAK/signal transducers and activators of the transcription (STAT) signaling pathway. This induces the expression of hundreds of ISGs that act antivirally. Among these, myxovirus resistance (Mx) proteins are key antiviral effectors that counter IAV infection (Haller et al., 2007). In humans, the Mx protein MxA represents the primary antiviral effector against IAV (H. Xiao et al., 2013). MxA is a key interferon-induced antiviral factor that restricts influenza virus replication by targeting viral ribonucleoprotein complexes (vRNPs). It has been proposed that it restricts influenza virus replication by targeting vRNPs potentially through interaction with nucleoprotein (NP). To suppress antiviral IFN signaling, IAV encodes the nonstructural protein 1 (NS1), a potent antagonist of the host cell IFN response (Hale et al., 2008; Ji et al., 2021). NS1 interferes with multiple steps of the antiviral signaling cascade. It binds to viral double-stranded RNA, which prevents recognition by pattern recognition receptors. This directly inhibits the sensors Retinoic Acid–Inducible Gene I (RIG-I) and Protein kinase R (PKR) and it also blocks host pre-mRNA processing. Collectively, these actions suppress the expression of antiviral ISGs (Dauber et al., 2006; Jureka et al., 2020; K. Zhang et al., 2019; L. Zhang et al., 2018)

The present study focuses on OP7, a distinct type of IAV DIP that carries several point mutations within segment 7 (Seg 7) of the IAV genome instead of a deletion (Dogra et al., 2023; Kupke et al., 2019). OP7 shows a higher antiviral activity than conventional IAV DIPs (Hein, Arora, et al., 2021; Hein, Kollmus, et al., 2021; Rand et al., 2021). So far, OP7 has primarily been investigated in Madin-Darby Canine Kidney (MDCK) cells. However, these cells lack a functional IFN response against IAV because their canine MDCK Mx proteins do not confer IFN-induced antiviral activity against human influenza strains (Seitz et al., 2010). To assess the relative contributions of the IFN response and the replication interference exerted by OP7, we used an IFN-competent human lung carcinoma cell line (Calu-3) in co-infection experiments (Felgenhauer et al., 2020; Hsu et al., 2012; Lokugamage et al., 2020). Overall, our results suggest that replication interference can support IFN induction, primarily by suppressing the viral IFN antagonist NS1. Furthermore, we demonstrate that OP7’s antiviral activity during low multiplicity of infection (MOI) STV co-infections is primarily mediated by stimulation of the IFN system. In addition, we tested the effective treatment window of OP7 in our *in vitro* model. We demonstrate that treatment with OP7 exerts antiviral effects even when cells are treated up to seven days before, and up to 24 h after STV infection. These findings suggest that OP7 is a promising antiviral candidate for further clinical testing. In addition, such mechanistic insights regarding the antiviral effects exerted by OP7 are essential to promote potential future regulatory and clinical acceptance of this novel class of antivirals.

## Materials and Methods

### Cells and viruses

Calu-3 cells (American Type Culture Collection (ATCC), #HTB-55) were maintained at 37 °C and 5 % CO_2_ in Dulbecco’s Modified Eagle Medium/Nutrient Mixture F-12 (DMEM/-F12) (Thermo-Fisher Scientific #11320033), supplemented with 10 % fetal bovine serum (Merck, #F7524), 1 % penicillin-streptomycin (Thermo-Fisher Scientific, #15140-122) and 1 % non-essential amino acids (Thermo-Fisher Scientific, #11140050). Influenza A/PR/8/34 H1N1 (PR8, provided by the Robert Koch Institute, #3138) was used as STV. MOI was calculated based on the tissue culture infectious dose (TCID₅₀) titer. For DIP infection, an OP7 “chimera” DIP was used that contains a large internal deletion in segment 1 (Seg 1) and point mutations in segment 7 of OP7 (Seg 7-OP7) (Dogra et al., 2023). PR8 and OP7 chimera DIPs were produced in a cell culture-based process in 125 mL shake flask, as described previously (Pelz et al., 2024). Subsequently, DIPs were purified by steric exclusion chromatography as described previously (Marichal-Gallardo et al., 2017; Pelz et al., 2024).

### Co-infection experiments

Calu-3 cells were seeded at a density of 1.5 × 10⁶ cells per well in 12-well plates and incubated for 24 hours. Next, the cells were washed once with phosphate-buffered saline (PBS) and infected with STV only at the indicated MOI or co-infected with highly concentrated OP7 chimera DIPs (2.67 E+11 particles per well) in a total volume of 600 µL. Cells were incubated for 1 h and washed once with PBS. Subsequently, 3 mL of DMEM/F12 was added. Where specified, cells were pretreated with ruxolitinib (ruxo) (Cayman Chemical; #11609) at a final concentration of 2 μM for 3 h prior to infection. After that, cells were washed once with PBS. During and post infection, ruxo at a final concentration of 2 μM was added to the media. At the indicated time points, samples were collected. Supernatants were clarified by centrifugation at 3000 × g for 5 min at 4 °C and aliquots of supernatants were stored at −80 °C until further virus quantification. For RNA extraction, the remaining cells were washed once with PBS, lysed with 350 µL RA1 buffer (Macherey-Nagel; #740955) and stored at −80 °C until RNA purification using the Nucleospin RNA isolation kit (Macherey-Nagel; #740955) according to the manufacturer’s instructions.

### Virus quantification

For the quantification of infectious viral particles, a TCID₅₀ assay was performed as described previously (Genzel et al., 2014). Total virus particle titers were quantified using a hemagglutination activity (HA) assay as described previously (Kalbfuss et al., 2008). HA titers were expressed as log_10_ hemagglutination units per 100 μL (log_10_ HAU/100 μL), allowing calculation of the total number of hemagglutination units in the virus preparation. The concentrations of DIPs (cDIP) were calculated from the hemagglutination (HA) titer using the equation below, where cRBC represents the red blood cell concentration (2.0 × 10⁷ cells/mL).

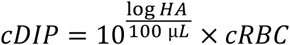

### Quantification of intracellular vRNA

Quantification of vRNA from purified intracellular total RNA was performed by real-time RT-qPCR as described previously (Kupke et al., 2019; Rand et al., 2021), but using 2× QuantiNova SYBR green qPCR master mix (Qiagen, #208056) for qPCR. The primers used for quantification of segment 5 (Seg 5) and segment 8 (Seg 8) are listed in (Kupke et al., 2019), and for Seg 7 and Seg 7-OP7 in (Hein, Kollmus, et al., 2021). RNA reference standards were used to allow for absolute quantification of individual vRNA segments.

### Gene expression analysis

mRNA expression levels in (co-)infected cells were quantified using real-time RT-qPCR as described previously (Kupke et al., 2019; Rand et al., 2021). Briefly, total RNA (500 ng) was reverse transcribed using an oligo(dT) primer and Maxima H Minus reverse transcriptase (Thermo Scientific, #EP0752) according to the manufacturer’s protocol. Next, qPCR was performed using gene-specific primers for MxA and IFN (Pelz et al., 2023) and 18S RNA as a housekeeping gene (18S for 5’-CGGACAGGATTGACAGATTG-3’, 18S rev 5’-CAAATCGCTCCACCAACTAA-3’). Gene expression was analyzed by the ΔΔCT method, normalized to 18S and expressed as fold-change relative to the mock infection control (untreated, uninfected cells).

### Quantification of viral proteins

Quantification of viral proteins was conducted as described previously (Küchler et al., 2022). In short, (co-)infected cells were washed once with PBS and then lysed with RIPA buffer (Thermo Fisher Scientific; #89900) and 26G needles (Terumo Agani; #AN2623R1). Subsequently, protein concentration was determined by a bicinchoninic acid assay (Thermo Fisher Scientific), followed by filter-aided sample preparation and proteolytic digestion. Stable isotope-labeled absolute quantification (AQUA) peptides were used for the absolute quantification of IAV peptides. Here, defined quantities of each AQUA peptide were spiked into the samples prior to analysis by Liquid Chromatography-Tandem Mass Spectrometry (LC-MS/MS). Quantification was performed by comparing the signal intensities of the endogenous (light) and isotopically labeled (heavy) peptides. Absolute copy numbers were calculated based on the known concentrations of the spiked standards.

## Results

### OP7 strongly suppresses IAV replication in human lung cells in vitro

To evaluate the antiviral effect of OP7 co-infection in IFN-competent human lung cells *in vitro*, Calu-3 cells were used. The cells were infected with STV alone at MOI 30, 3, or 10^-3^, or co-infected with OP7. At STV MOI 10^-^³, OP7 co-infection resulted in the complete suppression of total virus particle release (Fig. 1C), quantified by the HA assay. In contrast, for OP7 co-infections at STV MOI 3, only marginal suppression was observed (Fig. 1B). No suppression of total virus titers was observed in OP7 co-infections at STV MOI 30 compared to STV-only infection (Fig. 1A). However, infectious virus titers (quantified by the TCID_50_ assay) indicated a more pronounced antiviral effect. OP7 co-infections reduced the release of infectious virus particles by approximately one order of magnitude at STV MOI 30 at 72 hours post infection (hpi) (Fig. 1D) and by two orders of magnitude at STV MOI 3 (Fig. 1E). A complete suppression of the release of infectious virus particles was observed for OP7 co-infections at STV MOI 10^-^³ (Fig. 3F). These results demonstrate that OP7 markedly reduces infectious IAV production in Calu-3 cells *in vitro*, showing the strongest antiviral efficacy at low MOI conditions.

**Figure 1:**
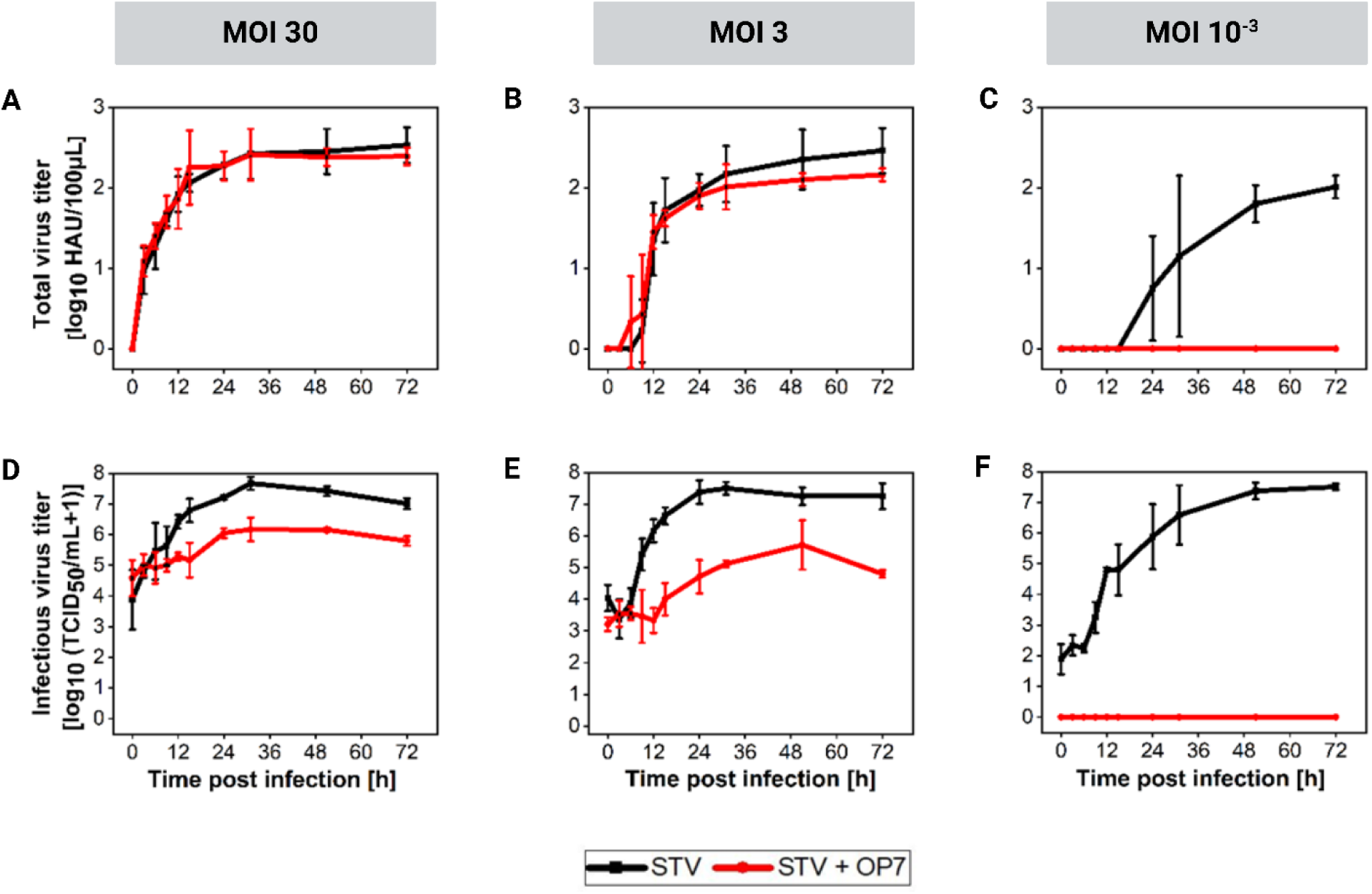
Inhibition of IAV replication by the defective interfering particle OP7 in human lung epithelial cells *in vitro*. Calu-3 cells were infected with STV alone (MOI 30, 3 and 10^-3^) or co-infected with OP7. Total virus titers were determined using HA assay (**A, B, C)**. Infectious virus titers were determined using TCID_50_ assay **(D, E, F)**. Error bars indicate the standard deviation of three independent experiments.

### OP7 exerts replication interference at high STV MOI co-infection

Next, we monitored replication interference to investigate its contribution to the antiviral effect exerted by OP7. For this, we quantified different vRNA segments during (co-)infections using real-time RT-qPCR. STV infection at MOI 3 resulted in efficient vRNA replication, with segments 5 and 7 steadily increasing until 12 hpi and reaching about 10^4^-10^5^ copies per cell at 48 hpi (Fig. 2A). In contrast, OP7 co-infection resulted in a distinct replication profile at STV MOI 3 (Fig. 2B). Seg 7-OP7 vRNA appeared to accumulated to higher levels, while the accumulation of other segments was reduced (∼10^3^-10^4^), consistent with previous observations (Rüdiger et al., 2024). This enhanced accumulation of Seg 7-OP7 vRNA and the concomitant decrease in STV vRNA replication indicate that OP7 co-infection causes antiviral replication interference (Rüdiger et al., 2024).

**Figure 2:**
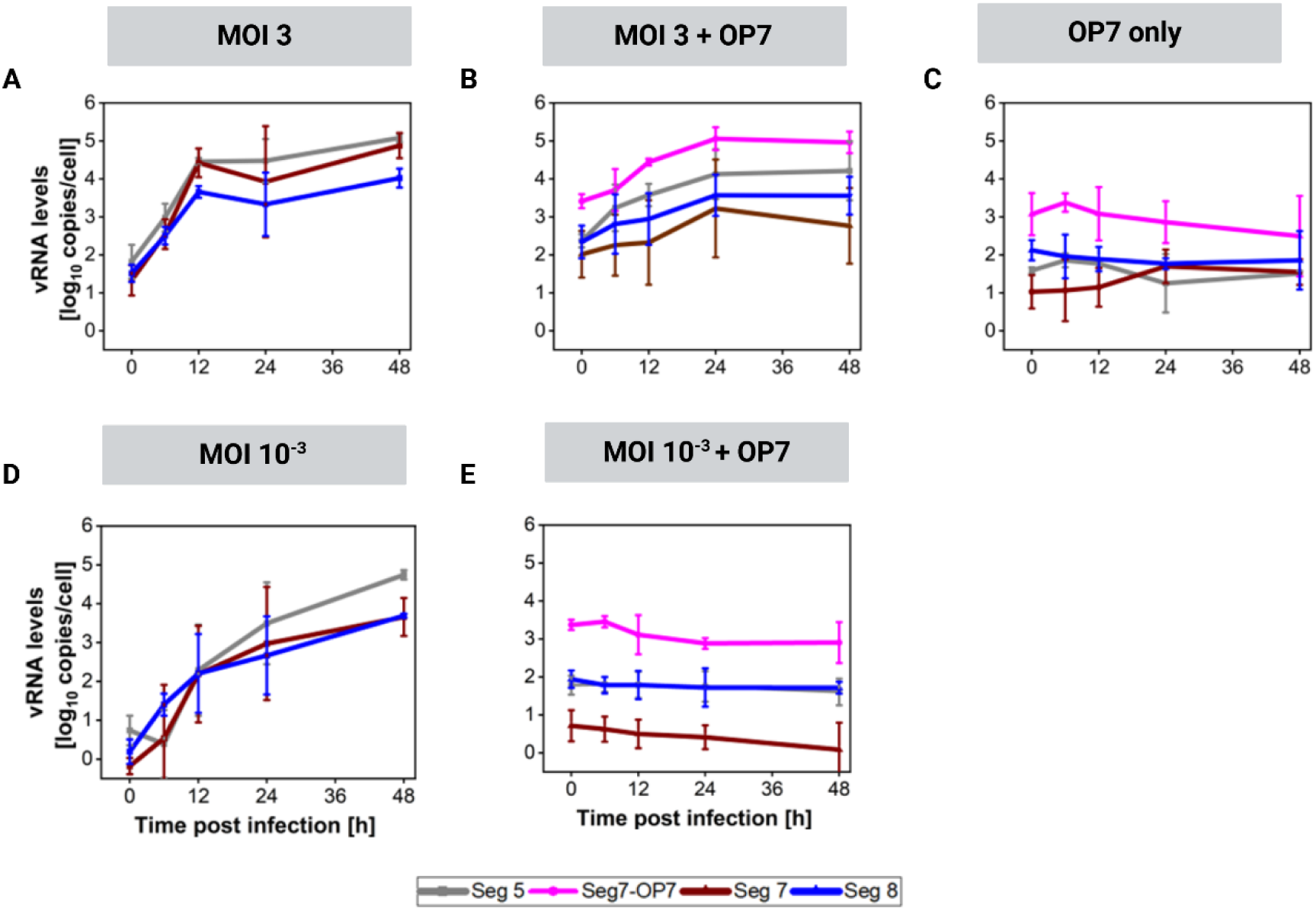
Intracellular genomic vRNA levels after OP7 co-infection. Calu-3 cells were infected with STV at MOI 3 or 10^-3^ (**A, D**), co-infected with OP7 at STV MOI 3 and 10^-3^ (**B, E**), or infected with OP7 only (**C**). Cells were lysed at indicated time points for subsequent intracellular RNA isolation, and the vRNA levels of Seg 5, Seg 7-OP7, Seg 7 and Seg 8 were quantified using real-time RT-qPCR. Error bars indicate the standard deviation of three independent experiments.

Surprisingly, no such replication interference could be observed for OP7 co-infections at STV MOI 10^-3^ (Fig. 2E). More specifically, there was no accumulation of Seg 7-OP7 vRNA or replication of other segments. For comparison, replication of vRNA progressed to high levels in STV-only infection at such low MOI (Fig. 2D). In addition, in an OP7-only infection, replication of all segments was absent, confirming that OP7 is a replication-incompetent virus particle (Fig. 2C). Taken together, pronounced replication interference was observed in co-infections with STV at a high MOI.

### Enhanced IFN signaling in OP7 co-infected cells is likely caused by suppression of the viral IFN antagonist NS1

Since complete suppression of replication was detected during low STV MOI co-infections with OP7 (Fig. 1C, F), while there was no evidence of replication interference (Fig. 2E), we next examined the stimulation of antiviral IFN induction under these conditions. For this, we quantified IFN-β1 and MxA expression by real-time RT-qPCR.

Co-infections with OP7 resulted in an earlier and enhanced expression of IFN-β1 and MxA compared to STV-only infections at MOI 3 (Fig. 3A, D) and MOI 10^-^³ (Fig. 3B, E). This effect was particularly pronounced at low MOI, where IFN-β1 and MxA expression levels exceeded those of STV-only infection by approximately two and one order of magnitude at 12 hpi, respectively. This enhanced cellular antiviral state may explain the antiviral effect exerted by OP7, particularly at low MOI co-infections. Similarly high expression of IFN-β1 and MxA was observed for OP7-only infection (Fig. 3C, F). This is consistent with the high intracellular quantity of vRNA (Fig. 2C) that likely leads to the observed early and enhanced stimulation of the IFN system. In comparison, decreased IFN signaling was observed for STV-only infections at high and low MOI (Fig. 3A, B, D, E), despite higher vRNA levels detected during the course of infection (Fig. 2A, D).

**Figure 3:**
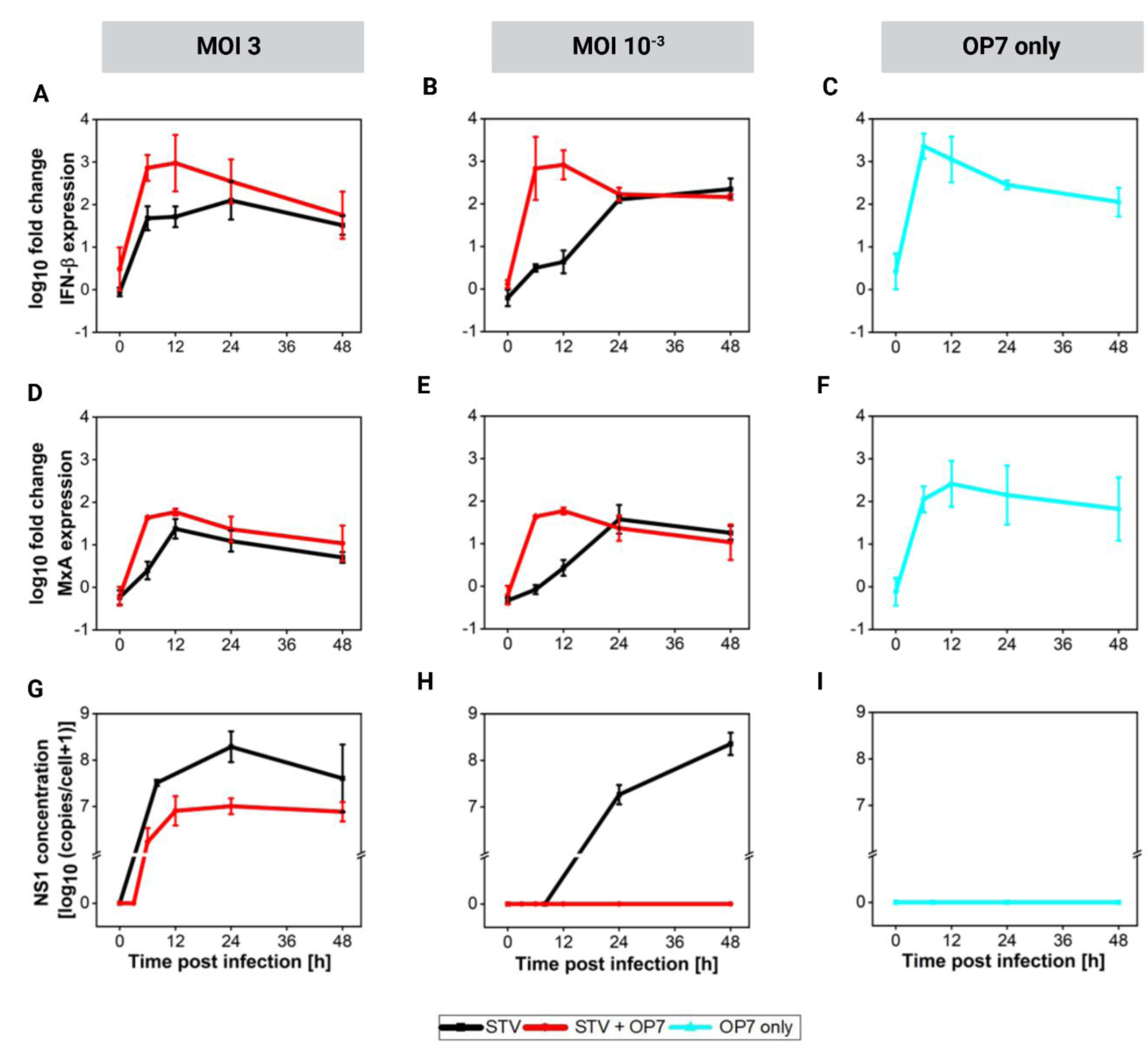
Induction of antiviral IFN-signaling through OP7 co-infections. Calu-3 cells were infected with STV at the indicated MOIs, co-infected with OP7, or infected only with OP7. Cells were lysed at indicated time points for subsequent intracellular RNA isolation and protein quantification. Gene expression of IFN-β (**A, B, C**) and MxA (**D, E, F**) was quantified using real-time RT-qPCR and expressed as fold change (relative to uninfected cells). Concentration of the viral NS1 protein was quantified via LC-MS/MS (**G, H, I**). Error bars indicate the standard deviation of three independent experiments.

To further investigate the cause of the elevated IFN response during OP7-only infection as well as OP7 co-infections, we quantified the viral IFN antagonist NS1 by LC-MS/MS. In STV-only infections at MOI 3, NS1 reached about 10⁸ copies per cell at 24 hpi (Fig. 3G). In contrast, for corresponding OP7 co-infection, the NS1 expression was reduced by more than one order of magnitude (Fig. 3G). OP7 exerts replication interference suppressing the accumulation of STV vRNAs, including that of Seg 8 (Fig. 2A, B), which encodes NS1. Reduced NS1 expression, in turn, likely leads to increased IFN responses during OP7 co-infection compared to STV-only infection at MOI 3 (Fig. 3A, D). Thus, replication interference appears to support OP7’s ability to stimulate enhanced IFN signaling at high STV MOI.

In both OP7 co-infection at STV MOI 10^-^³ and OP7-only infection, the NS1 protein was undetectable (Fig. 3H, I), likely due to the low quantity of Seg 8 (Fig. 2C, E). This may explain the elevated IFN response under these conditions (Fig. 3B, C, E, F). Taken together, these results demonstrate that OP7 strongly stimulates antiviral IFN signaling, at least in part through suppression of the IFN antagonist NS1. This enhances the antiviral IFN signaling in co-infected cells. In addition, IFN induction appears to explain most of the antiviral effect of OP7 observed at low STV MOI co-infection conditions.

### Antiviral activity of OP7 at low STV MOI co-infection appears to be primarily driven by IFN stimulation

In order to investigate the contribution of IFN stimulation to the antiviral effect of OP7 at low STV MOI conditions, we next used the small-molecule drug ruxo to reduce IFN signaling (Pelz et al., 2023; Rand et al., 2021). We observed an inhibition of MxA expression in the presence of ruxo (Fig. 4B). However, it appears that the residual MxA expression observed under these conditions still resulted in a complete inhibition of infectious virus particle release (Fig. 4A). The inhibition does not appear to be caused by replication interference, as vRNA accumulation was absent (Fig. 4C-E).

**Figure 4:**
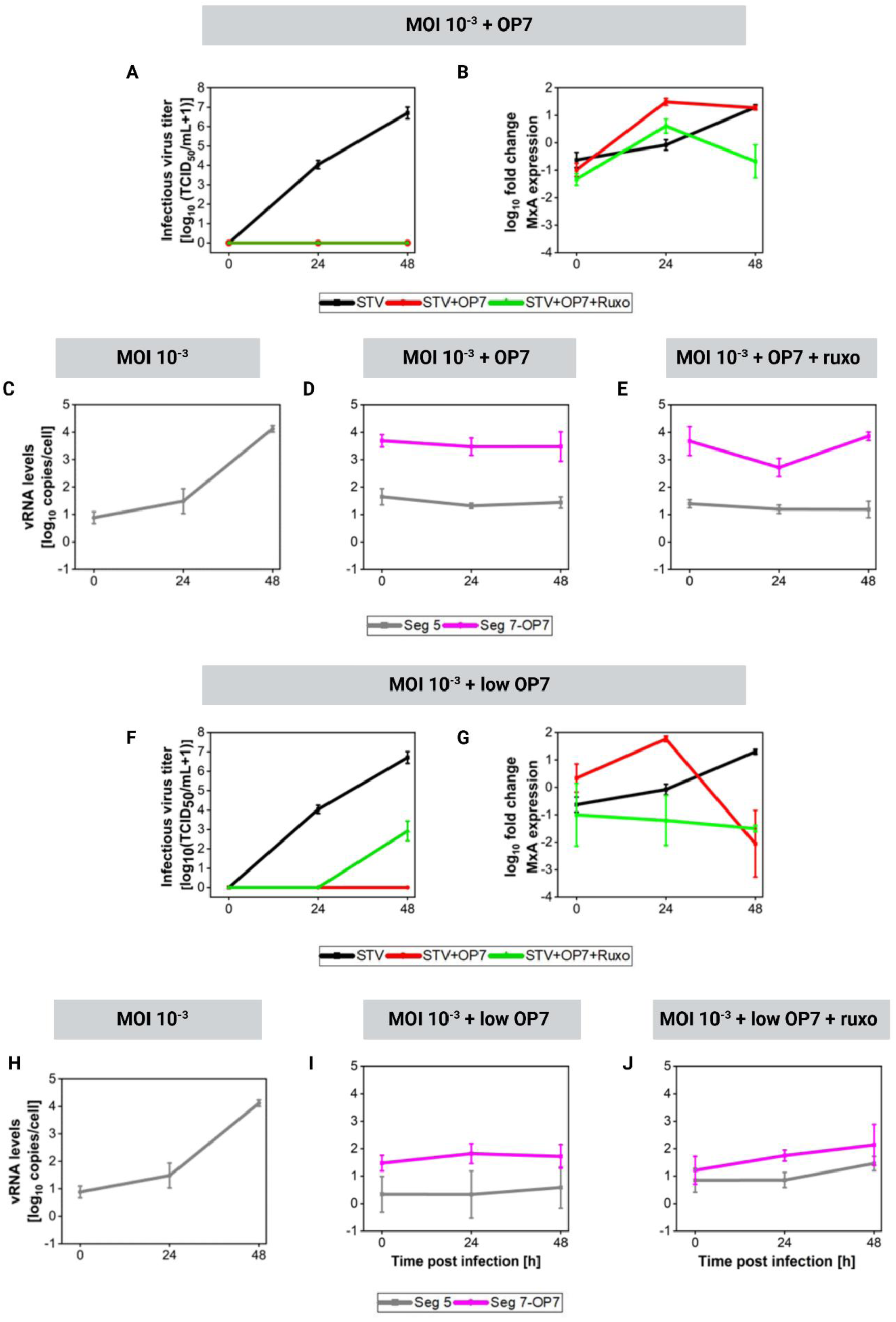
Impact of Mx expression on the antiviral effect at low STV MOI. Calu-3 cells were infected with STV at MOI 10^-3^ or co-infected with OP7 or a 100-fold dilution of OP7 (“low OP7”). Where specified, cells were pretreated with ruxo at a final concentration of 2 μM for 3 h prior to infection. Supernatants were taken at indicated time points and infectious virus titers were quantified using the TCID_50_ assay (**A, F**). Cells were lysed at indicated time points for subsequent intracellular RNA isolation. MxA was quantified using real-time RT-qPCR (**B, G**) and expressed as fold change (relative to uninfected cells). Seg 5 and Seg 7-OP7 vRNA levels were quantified using real-time RT-qPCR (**C, D, E, H, I, J**). Error bars indicate the standard deviation of three independent experiments.

Next, we reduced the OP7 dose by 100-fold (“low OP7”) during co-infection. Surprisingly, OP7 still fully suppressed infectious virus titers at STV MOI 10^-3^ in the absence of ruxo (Fig. 4F), which emphasizes the strong antiviral effect exerted by OP7. When ruxo was present, MxA expression was inhibited to a level slightly below that of basal gene expression (Fig. 4G). Under these conditions, the inhibitory potential of OP7 was reduced, as evidenced by a reduced suppression of infectious virus particle release compared to the co-infection without ruxo (Fig. 4F). The residual inhibition observed in the presence of ruxo may be explained by marginal replication interference (Fig. 4H-J). Taken together, the antiviral activity of OP7 at low STV MOI appears to be primarily caused by the stimulation of the IFN system.

### Antiviral activity upon OP7 infection for up to seven days before and 24 h after STV infection in human lung cells in vitro

Next, we tested the applicability of OP7 as an antiviral agent, by investigating OP7 infection up to 14 days before and up to 48 h after STV infection. This corresponds to prophylactic and therapeutic treatment scenarios, respectively.

In order to assess the prophylactic antiviral effect of OP7 *in vitro*, we treated cells with OP7 and subsequently challenged with STV (MOI 3 or MOI 10^-3^) at 3, 7, 10, and 14 days post treatment. We observed an antiviral effect for OP7 pre-treatment for up to seven days (Fig 5A, B). Specifically, at STV MOI 3, OP7 pre-treatment reduced infectious virus particle release by up to three orders of magnitude when challenged with STV simultaneously or three days after OP7 treatment (Fig. 5A). After seven days of pre-treatment with OP7, a marginal suppression was observed when challenged with STV MOI 3. At STV MOI 10^-3^, OP7 treatment led to a complete suppression of infectious virus particle release when applied simultaneously (Fig. 5B). Reduction of infectious viral titers comprised over three, or one order of magnitude, when challenged with STV three or seven days after OP7 application, respectively. In addition, we monitored Seg 7-OP7 vRNA levels and MxA expression in OP7-only treated cells over time. Seg 7-OP7 vRNA could be detected for up to 14 days, indicating high vRNA stability in cells (Fig. 5C). In addition, we detected a long-lived and enhanced induction of innate immunity, as indicated by MxA expression in OP7-only infected cells compared to mock-infected cells (Fig. 5F). The loss of protection observed at ten days post infection might be explained by the continual decline in Seg 7-OP7 vRNA levels and MxA expression over time.

**Figure 5:**
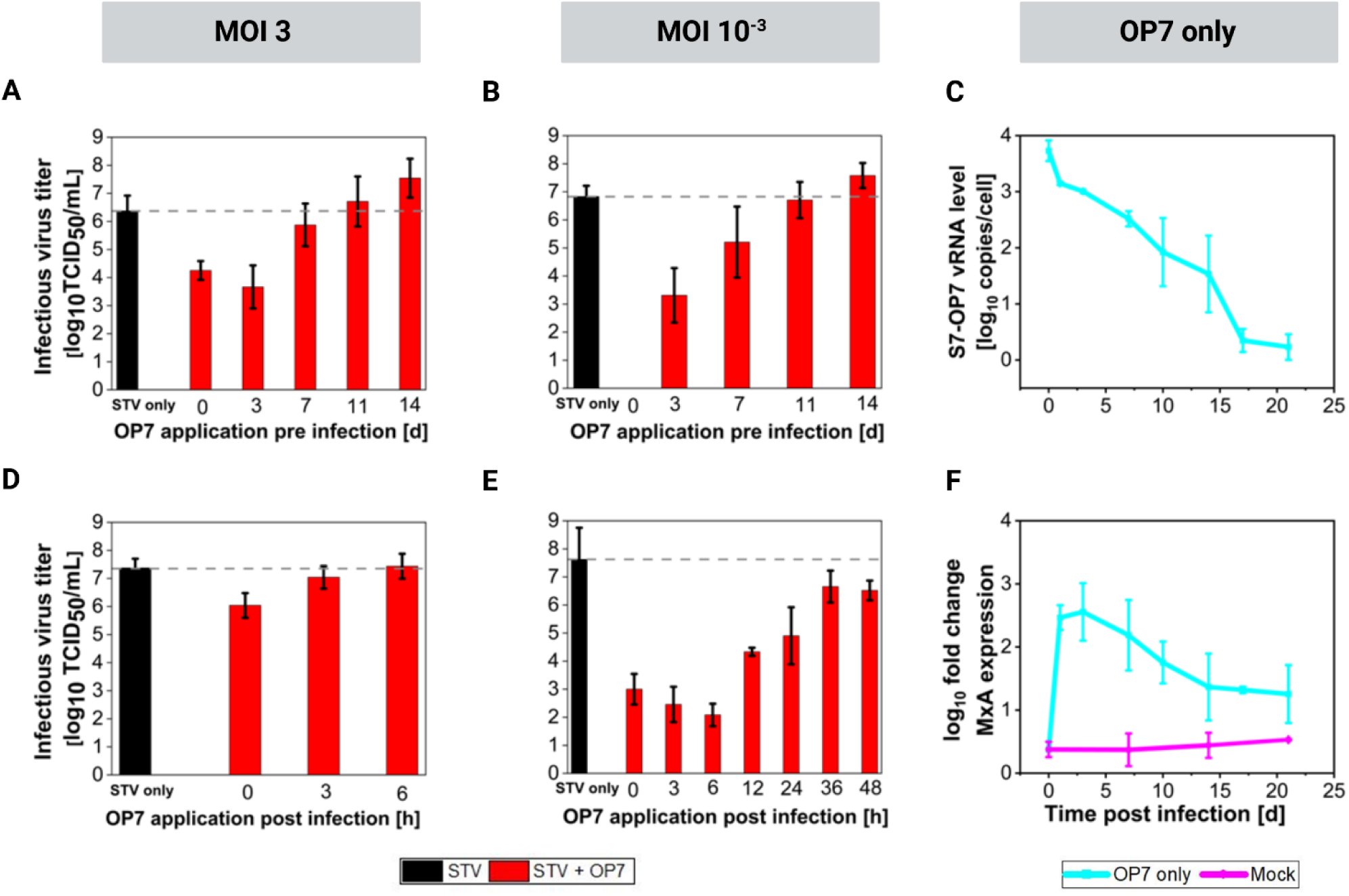
Prophylactic and therapeutic antiviral efficacy of OP7 in human lung epithelial cells *in vitro*. Calu-3 cells were infected with OP7 at indicated time points before (**A-B**) or after STV infection (**D-E**) at indicated MOIs. Infectious virus titers were determined using the TCID_50_ assay at 72 hpi (**A-B**) or 48 hpi (**D-E**). The dashed lines indicate infectious virus titers of STV only infections. OP7-only infected cells were lysed at indicated time points for subsequent intracellular RNA isolation. Seg 7 OP7 vRNA levels were quantified using real-time RT-qPCR (**C**). MxA (of OP7-only and mock-infected cells) was quantified using real-time RT-qPCR and expressed as fold change (relative to uninfected cells) (**F**). Error bars indicate the standard deviation of three independent experiments.

To assess the therapeutic effect of OP7 treatment, we infected Calu-3 cells with STV at MOI 3 or MOI 10^-3^ and subsequently treated cells with OP7 up to 48 hpi (Fig. 5D, E). At STV MOI 3, we detected a weak reduction in viral replication when OP7 was applied for up to three hours post infection (Fig. 5D). At STV MOI 10^-3^, OP7 addition resulted in significant suppression of infectious viral titers by up to five orders of magnitude when applied at 0, 3 or 6 hpi, and up to three logs when applied at 12 and 24 hpi (Fig. 5E). This therapeutic antiviral effect appeared to be lost, when OP7 was applied at 36 or 48 hpi. Overall, these findings demonstrate that OP7 confers a robust and long-lasting prophylactic antiviral activity for up to seven days and has substantial therapeutic efficacy when administered early after infection, particularly under low MOI conditions. This highlights its potential as a preventive and post-exposure antiviral agent *in vitro*.

## Discussion

DIPs have been suggested as promising, broad-spectrum antivirals. We investigated the antiviral effects of OP7 *in vitro*, an unconventional IAV DIP with a unique genetic structure. Our results demonstrate that OP7 efficiently suppresses the replication of STV vRNA genome segments through replication interference. In addition, this inhibition appears to enhance IFN stimulation by suppressing the expression of the viral IFN antagonists NS1. Furthermore, IFN stimulation appears to contribute largely to the antiviral effect of OP7 observed under low STV MOI co-infection conditions, where it completely abrogates viral replication. Finally, we show that OP7 infection results in antiviral activity that lasted for up to seven days, corresponding to a possible prophylactic treatment window. In addition, OP7 co-infection for up to 24 h after STV infection still showed pronounced antiviral effects (therapeutic setting). These results clearly suggest its applicability as an antiviral.

Our data demonstrate that OP7 co-infection potently suppresses the release of infectious IAV, particularly at low STV MOI. Even at low (100-fold diluted) OP7 concentrations, complete suppression of virus replication was observed (Fig. 4F), demonstrating its strong antiviral potential. The natural infectious dose for airborne IAV transmission in humans has been estimated to be approximately three TCID₅₀ units (Alford et al., 1966), corresponding to a low MOI infection scenario. In conclusion, OP7’s strong antiviral activity during low STV MOI co-infection further supports its potential as a potent antiviral agent (Kupke et al., 2019; Hein, Kollmus, et al., 2021; Rand et al., 2021; Dogra et al., 2023; Pelz et al., 2023, 2024; Rüdiger et al., 2024).

The higher antiviral effects of OP7 at low STV MOI may be explained by paracrine IFN signaling. Released IFNs from OP7 (co-)infected cells can act on neighboring, uninfected cells, inducing an antiviral state early, before STV infection occurs, thereby suppressing or preventing STV replication, a mechanism that was also proposed for conventional IAV DIP (Russell et al., 2019). In contrast, at high STV MOI infections (e.g., MOI 3), nearly all cells become infected simultaneously. Accordingly, viral replication may proceed too rapidly for IFN-mediated autocrine and paracrine signaling to suppress it. Therefore, in high STV MOI co-infection scenarios with OP7, replication interference appears to play a more crucial role in inhibiting STV replication.

Quantification of vRNA at high STV MOI co-infections revealed enhanced accumulation of the mutated Seg 7-OP7 vRNA, while the replication of STV vRNA segments was strongly suppressed. This indicates that OP7 interferes directly with viral replication, likely by competing for viral polymerase complexes and nucleoproteins required for S7-OP7 vRNA synthesis, as previously demonstrated in a systems virology approach (Rüdiger et al., 2024). These findings align with the established concept of replication interference mediated by DI genomes that outcompete STV genomes for viral proteins, such as polymerases, thereby limiting STV propagation (Akkina et al., 1984; Perrault, 1981; Widjaja et al., 2012; Wu et al., 2022). Importantly, replication interference is not limited to DIPs of IAV. It has also been reported for DIPs derived from a diversity of viruses, including SARS-CoV-2, poliovirus, Nipah virus, Zika virus, HIV, and others (Chaturvedi et al., 2021; Pitchai et al., 2024; Rezelj et al., 2021; Sharov et al., 2021; Shirogane et al., 2021; Vignuzzi & López, 2019; Welch et al., 2022; Y. Xiao et al., 2021). The source of the replicative advantage differs among the DIP’s virus species and may result from enhanced polymerase binding, improved encapsidation, or increased replication efficiency due to specific genome rearrangements or mutations (Calain & Roux, 1995; Grabau & Holland, 1982; Kupke et al., 2019; Perrault, 1981). The enhanced replication of Seg 7-OP7 vRNA in OP7 co-infections can be explained by the presence of a “superpromoter” on Seg 7-OP7 vRNA (Belicha-Villanueva et al., 2012; Kupke et al., 2019; Rüdiger et al., 2024; Vreede et al., 2008).

Interestingly, suppressing STV vRNA replication during co-infections with OP7 at high STV MOI also reduces expression of the viral NS1 protein, a known antagonist of IFN signaling (Kochs et al., 2007). This reduced expression may explain the enhanced IFN and Mx stimulation observed during OP7 co-infections, which results in an enhanced antiviral state. This concept of a decrease in the expression of virus-encoded antagonists of innate immunity, mediated by replication interference and leading to a stronger antiviral state in DIP co-infected cells, was also suggested elsewhere (Genoyer & López, 2019). Thus, OP7 appears to promote antiviral effects by directly suppressing STV genome segment replication and indirectly by reducing the expression of virus-encoded IFN antagonists. Thus, the antiviral effect of OP7 arises from a synergistic interplay between replication interference and IFN activation; the relative importance of each depending on the infection dynamics. While replication interference seems to be the dominant antiviral mechanism at high STV MOI co-infections, IFN activation appears to largely contribute to inhibition during low STV MOI conditions.

For OP7 infection in Calu-3 cells *in vitro*, we observed a prophylactic and therapeutic treatment window of up to seven days and 24 h, respectively, where a pronounced suppression of STV replication was evident. Therefore, we can confirm a long treatment window for the IAV DIP OP7, which was previously also reported for conventional IAV DIPs *in vivo* (Dimmock et al., 2008). More specifically, mice receiving a dose of conventional IAV DIPs seven days prior to a lethal IAV challenge were fully protected from death and clinical disease (Dimmock et al., 2008). In another study, the intratracheal administration of IAV DI-vRNA-encoding plasmids into mice led to antiviral effects for up to three days (Zhao et al., 2018, 2022). Similarly, poliovirus DI genomes delivered intranasally by lipid nanoparticles (LNPs) two days before lethal poliovirus infection protected 90% of mice from death (Y. Xiao et al., 2021). With respect to the therapeutic potential of DIPs, administration of conventional IAV DIPs protected mice from a lethal IAV dose even when administered one day post infection and disease was delayed when administered two days post infection (Dimmock et al., 2008). Intratracheal administration of plasmids encoding IAV DI-vRNA up to 24 hours after a lethal viral challenge resulted in markedly improved survival rates of up to 90%, depending on the challenge IAV strain (Zhao et al., 2018). Similarly, SARS-CoV-2 DI genomes in LNPs administered 12 hpi after SARS-CoV-2 infection significantly reduced viral load and lung pathology in hamsters (Chaturvedi et al., 2021). In the context of poliovirus infection, the therapeutic administration of poliovirus DI-vRNA in LNPs up to two days after a lethal challenge conferred substantial protection and yielded a 70% survival rate in mice (Y. Xiao et al., 2021).

Small-molecule antivirals to treat IAV infections, which are the current treatment standard in the clinic, are the neuraminidase inhibitors zanamivir and oseltamivir (Gillissen & Höffken, 2002; Jefferson et al., 2014). Of note, a direct comparison of administration of both antivirals or conventional IAV DIPs showed enhanced antiviral activity of the DIPs, in both prophylactic and therapeutic treatment scenarios (Dimmock et al., 2012; Zhao et al., 2018, 2022). The pronounced antiviral activity observed when OP7 is applied early suggests that DIPs must be present during or shortly after infection to outcompete the standard virus for replication machinery. Thus, similar to small-molecule drugs, early administration is critical for optimal therapeutic benefit, but OP7 may offer an additional advantage by simultaneously causing the expression of ISGs. In this context, the broad-spectrum antiviral activity of IAV DIPs (mediated by IFN stimulation), e.g. against replication of other clinically relevant respiratory viruses, including influenza B virus, SARS-CoV-2, and respiratory syncytial virus (Pelz et al., 2023; Rand et al., 2021; Scott et al., 2011a), would be highly attractive in pandemic scenarios. Their use could help to contain the spreading of newly emerging IFN-sensitive viruses, especially in scenarios when vaccines or antivirals are not available yet. Finally, the development of resistance of viruses to the DIP’s antiviral effects is regarded very unlikely according to current research (Chaturvedi et al., 2021; Pitchai et al., 2024; Rast et al., 2016). For example, no resistant IAV strains have been reported to date. This demonstrates the potential of the use of DIPs as robust antiviral agents that offer clear advantages compared to small-molecule antivirals, which are susceptible to viral resistance development (Hedskog et al., 2023; Lu et al., 2021)

Future studies on the antiviral activity of OP7 *in vitro* may include 2D and 3D tissue models under culture conditions that more closely mimic the mucosal environment of the human airway. Combined with mathematical modeling, these approaches could support the optimization of OP7 dosing and treatment timing strategies. This would have implications for designing clinical trials for OP7 as a new antiviral treatment modality.

## Author contributions

Conceptualization, P.O., S.Y.K. and U.R.; formal analysis, P.O., J.K.; investigation, P.O., J.K, K.H., E.H.; writing—original draft preparation, P.O.; writing—review and editing P.O., D.R., J.K, S.Y.K. and U.R.; visualization, P.O.; supervision, S.Y.K. and U.R.; project administration, P.O. and S.Y.K. All authors have read and agreed to the published version of the manuscript.

## Acknowledgements

We thank Nancy Wynserski and Claudia Best for their excellent technical assistance.

